# Discovery of Highly Active Kynureninases for Cancer Immunotherapy through Protein Language Model

**DOI:** 10.1101/2024.01.16.575968

**Authors:** Hyunuk Eom, Kye Soo Cho, Jihyeon Lee, Stephanie Kim, Sukhwan Park, Hyunbin Kim, Jinsol Yang, Young-Hyun Han, Juyong Lee, Chaok Seok, Myeong Sup Lee, Woon Ju Song, Martin Steinegger

## Abstract

Overcoming the immunosuppressive tumor microenvironment is a promising strategy in anticancer therapy. L-kynurenine, a strong immunosuppressive metabolite can be degraded through kynureninases. Through homology searches and protein language models, we identified and then experimentally determined the efficacy of four top-ranked kynureninases. The catalytically most active one nearly doubles turnover number over the prior best, reducing tumor weight by 3.42 times in mouse model comparisons, and thus, presenting substantial therapeutic potential.

---

Popular cancer immunotherapies include the administration of monoclonal antibodies to block immune checkpoints (ICPs) like PD-1 and PD-L1 and the adoptive transfer of T cells targeting cancer cells^1^. However, the objective response rate for solid tumors is still low (< 30%), necessitating the development of additional choices^1^.

Tumor microenvironments (TME) are known to be immunosuppressive, negating the functions of diverse immune cells including primary tumor-attacking cells like CD8^+^ T and NK cells, thus promoting the tumor growth^2^. In addition to ICPs, tumors contain other strong immunosuppressive factors/mediators including L-kynurenine (L-KYN)^3^. L-KYN as an AhR ligand induces T cell apoptosis, Treg induction, and PD1 upregulation^3,4^. Thus, removing the L-KYN in the TME could be a good therapeutic strategy.

Indeed, Triplett et al. previously demonstrated that metabolizing L-KYN with exogeneous kynureninases (KYNase) can enhance the immune response against cancer^5^. In particular, KYNase from *Pseudomonas fluorescens* (Pf-K) shows the high catalytic efficiency (*k*_cat_/*K*_M_) compared to the human-encoded KYNase (K0). The administration of Pf-K in combinations with ICP inhibitors or a cancer vaccine boosted the active CD8^+^ T cell numbers, reduced the tumor growth, and increased the mouse survival time by 45%^5^.

Since Pf-K showed a good therapeutic benefit in mouse model, it is worth of exploring a broader range of KYNases that might offer even greater catalytic efficiency and therapeutic potential. One way to detect more efficient enzymes is by searching for homologous proteins in a large protein database^6^ and checking their activity experimentally. However, prioritizing the experiments to execute, while facing thousands of potential homologous candidates remains a major challenge. Large protein language models (pLLMs), pre-trained on vast datasets ranging from millions to billions of amino acid sequences, have acquired knowledge about both protein function and structure.^7^ Consequently, they offer a solid foundation to developing predictive models, even with only sparse experimental data points available.

To find candidate KYNases, we propose to utilize homology search and pLLMs^7^ to find and rank the catalytic activities of KYNases (Fig. 1a). we searched with MMseqs2^6^ for sequences similar to Pf-K, across database containing 3.2 billion protein sequences^8-10^ extracted from known species and uncultivated sources. We identified 10,195 sequences that closely match Pf-K over at least 90% of their lengths and meet a strict E-value threshold of less than 0.001. To remove redundant sequences, we grouped them based on 90% sequence identity, resulting in a set of 5,690 sequences. We then used MMseqs2 taxonomy^6^ to assign a likely species origin to each sequence. The taxonomical distribution of these sequences across different species is shown in Supplementary Fig. 1.

**Figure 1.**
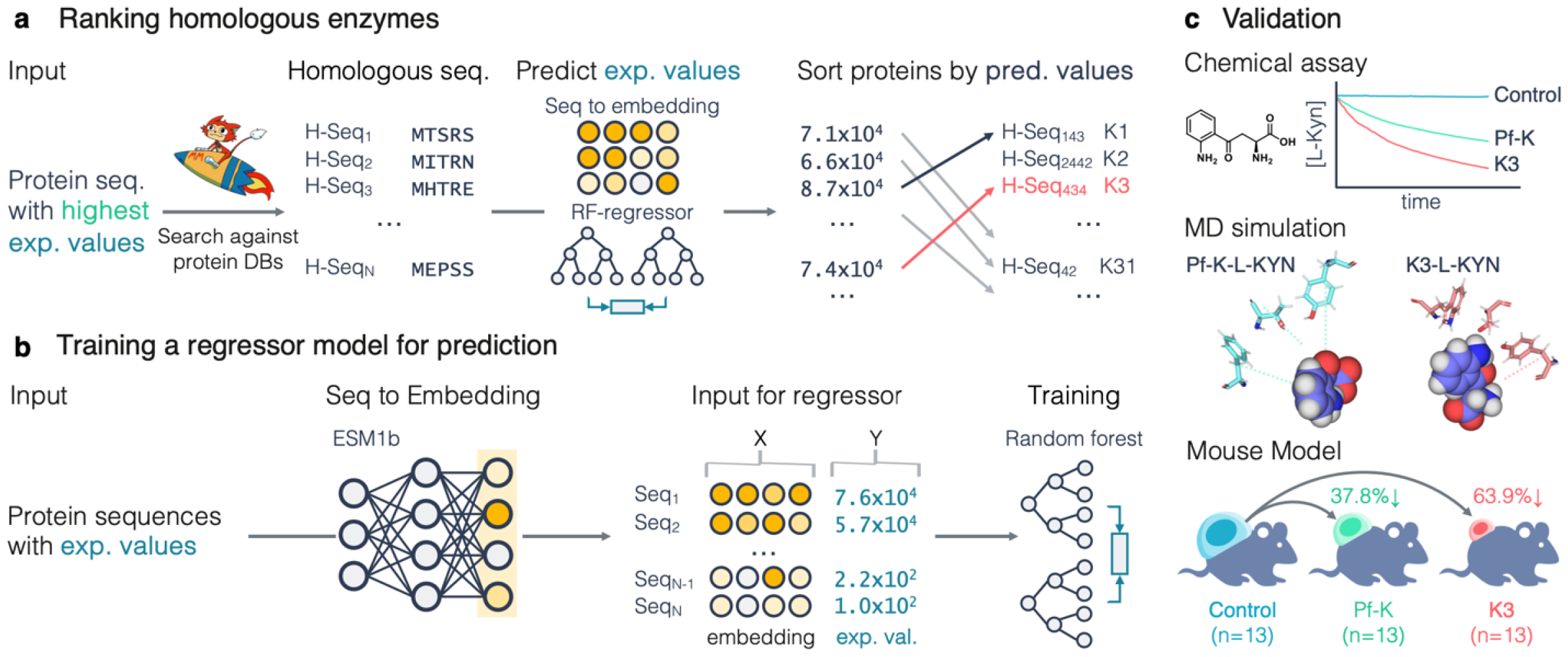
Workflow illustrating the discovery and validation of novel kynureninases for improving cancer immunotherapy. **a**, Search for homologs of the most catalytically active KYNases (Pf-K), embed them, and rank the hits based on their predicted activity. **b**, Training a regressor model to predict the enzymatic activity from protein using experimentally determined values and pLLM embeddings (indicated in yellow). **c**, Validate the four top-ranked homologs in three ways. Catalytic activity assay for monitoring the consumption of L-KYN: the chemical structure and time-dependent consumption of L-KYN. Molecular dynamics (MD) simulations demonstrating the interactions of L-KYN and KYNases. Mouse experiments, depicting the antitumor activity of KYNases.

For each detected protein sequence, we predicted the kinetic parameters of catalytic efficiency (*k*_cat_/*K*_M_), using a predictor based on embeddings generated by the ESM pre-trained language model^7^. Our predictor transforms protein sequences into fixed-size vector-space embeddings using language models, then predicts *k*_cat_/*K*_M_ via a random forest regression model. Trained on an 80/20 split of 159 experimentally measured sequences (Fig. 1b), this model achieves a Spearman correlation of 0.813, denoting high predictive accuracy^5,11^ (Fig. 2a). We implemented an easy-to-use webserver (see Methods and Data availability) for researchers to conduct similar analyses using their own measures.

**Figure 2.**
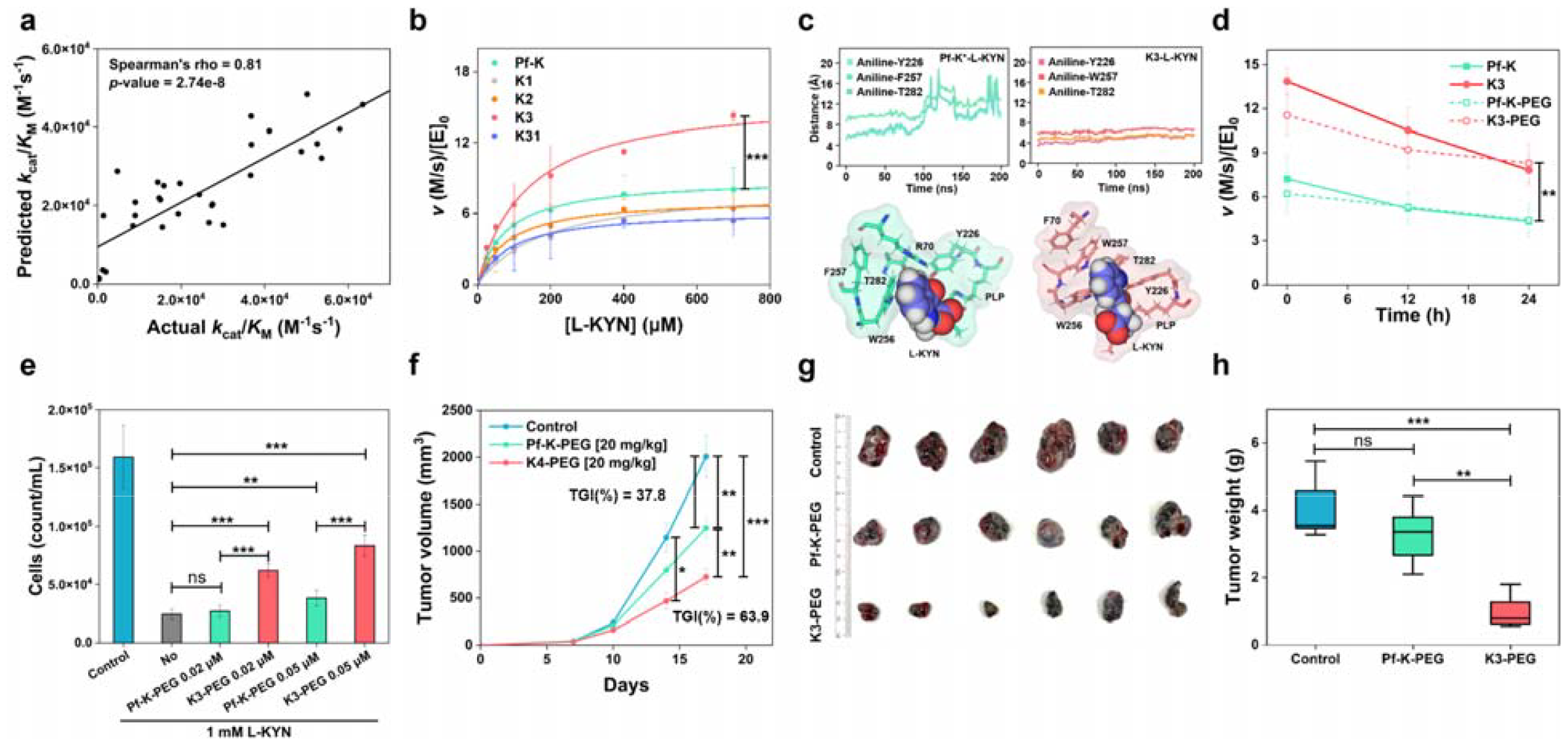
Computational and experimental results. **a**, Correlation plot between predictions and experimentally determined *k*_cat_/*K*_M_ values on our trains set. **b**, Michaelis-Menten kinetic analysis **c**, Average atomic distances over simulation time (200 ns) between N of aniline ring of L-KYN and corresponding residues in Pf-K^*^ and K3. Snapshots at the end of simulation for Pf-K* and K3. **d**, Catalytic activities after incubation at 37 °C to measure catalytic stability. **e**, *In vitro* T cell activation/proliferation under L-KYN. CD8^+^ T cell counts measured after T cell stimulation (at day 7). **f**, Tumor growth curve of B16-F10 bearing mice (C57BL/6) that were treated with peritumoral injection of control (PBS, n=13), Pf-K-PEG (20 mg/kg, n=13) and K3-PEG (20 mg/kg, n=13). **g**, Representative tumor images and **h**, tumor weight after isolation at the end of monitoring. Statistical significance was verified for t-test (two-tailed), *p* value (^*^ *p* ≤ 0.05, ^**^ *p* ≤ 0.01, ^***^ *p* ≤ 0.001). Error bars represents the mean ± SEM.

We ranked the 5,690 protein sequences detected by MMseqs2 using our RF predictor. The 10 highest ranked sequences were all prokaryotic, 4 were predicted to be from the *Pseudomonas* genus and the remaining 6 were from various *Proteobacteria*. As a general trend, our predictor predicts higher *k*_cat_/*K*_M_ for sequences from bacteria than from eukaryotes. Next, we chose four high-ranking sequences for validation. The highest-ranking was a KYNase from the *Pseudomonas* genus that has a 70.6% sequence identity to K-Pf; the second best was from *Pseudomonas* sp. (K2), and third *Bordetella genomosp*. (K3) with a 64.42% sequence identity. For further comparison, we selected an additional sequence from *Rheinheimera* sp., ranked 31^st^ (K31), for heterologous expression in *E. coli* (Supplementary Fig. 2 and Supplementary Table 1).

All four putative KYNases we selected show comparable or even substantially higher kinetic parameters than Pf-K^5,12^ (Fig. 2b, Supplementary Fig. 3, and Supplementary Table 2); in particular, K3 exhibits a ∼2-fold higher turnover number (*k*_cat_) while *k*_cat_/*K*_M_ value is comparable with that of Pf-K (Fig. 2b and Supplementary Figs. 3 and 4). Despite the inconsistency between the predicted order in *k*_cat_/*K*_M_ and the experimental results, these findings underscore the effectiveness of our predictor in identifying enzymes with high catalytic efficiency. Furthermore, these results also indicate that K3 may be a promising candidate for therapeutic applications.

To understand the molecular origin of the high catalytic activities of K3, we conducted molecular dynamics simulations^13^ (Supplementary Figs. 5 and 6). K3 exhibits improved binding of L-KYN (Fig. 2c), possibly because W257, Y226, and T282, form a tightly packed active site and leads to more stable substrate-enzyme interactions. In contrast, the corresponding residues in K0 and Pf-K are Y226, F257, and T282, forming a less compact binding site with increased conformation flexibility. Furthermore, the binding site of K3 is more packed that that of Pf-K; its cavity volume is 0.3−0.8 Å^3^ smaller than Pf-K (Supplementary Table 3).

The pH-dependent activity and thermal stability of enzymes can be critical in cancer treatments^14^ (Supplementary Fig. 7). Although K3 shows lower catalytic activity at acidic pH conditions found in tumors, it retained sufficiently higher activities relative to Pf-K. The thermal stability of K3 was comparable with that of Pf-K; the melting temperature (*T*_m_) of Pf-K and K3 were 60.8 °C and 61.1 °C, respectively. PEGylation negligibly altered the activity (86% and 83% relative to their non-PEGylated ones) and melting temperature (57.4 °C and 60.0 °C) (Fig. 2d and Supplementary Fig. 8). In addition, K3 was superior to Pf-K in the reaction with L-KYN measured after 24 h incubation at 37 °C, implying sufficient stability and activity for *in vivo* applications (Fig. 2d).

To investigate whether Pf-K-PEG and K3-PEG can reverse L-KYN -induced inhibition of the activation/proliferation of CD8^+^ T cells, Pf-K-PEG and K3-PEG were treated in the presence of L-KYN. The size of T cell groups formed after T cell proliferation was reversed more strongly in the culture treated with K3-PEG than Pf-K-PEG (Supplementary Fig. 9). In addition, the cell counts showed the similar pattern to the aggregate T cell group size (Fig. 2e) (*p*-value=0.039, on day 7). This result indicates that K3-PEG is functionally more active than Pf-K-PEG in the biologically relevant context.

To evaluate the anti-tumor efficacy of Pf-K-PEG and K3-PEG *in vivo*, we employed a syngeneic mouse melanoma implant tumor model and administrated PEG-KYNase peritumorally as previously reported^5^. As expected, K3-PEG-treated group showed much stronger tumor growth inhibition (TGI(%)=63.9, p-value=0.00008) than that of Pf-K-PEG (TGI(%)=37.8, p-value=0.0067) (Fig. 2f-h and Supplementary Fig. 10). However, the KYNase-treated mice showed no significant body weight change compared to the control (Supplementary Fig. 11). Additionally, in vivo anti-tumor efficacy test using colon cancer mouse model showed similar result (Supplementary Fig. 12). These results together indicate that K3-PEG is an effective anti-cancer enzyme, validating our enzyme search algorism.

Trp is rapidly degraded into L-KYN by IDO1 or TDO in tumors. The resulting Trp depletion and/or L-KYN formation make a strong immunosuppressive TME^1,2,3,4^. Accordingly, multiple IDO1 inhibitors have been developed. Although the most advanced, highly specific IDO1 inhibitor, epacadostat, showed encouraging results in a phase I/II trial in combination with anti-PD1 Ab, a large phase III trial was unsuccessful^15^, possibly because it cannot inhibit other Trp degrading enzymes, such as TDO and IDO2^3,15^. KYNases have several pharmacological advantages over the small molecule IDO/TDO inhibitors. First, complete blockade of Trp catabolism by IDO1 and TDO dual inhibitors may induce serious side effects such as high level of Trp-induced neurological effects and blockade of synthesis of downstream neuroprotective compounds^5^. In contrast, KYNase may not greatly perturb the Trp catabolism by removing the metabolites in the downstream. Secondly, small molecule inhibitors of enzymes frequently result in the acquisition of drug-resistant mutations^16^, while KYNase is not relevant for such mechanism since it degrades the immunosuppressive metabolite itself. Our work demonstrated that homology search combined with pLLMs can detect and prioritize highly catalytically active therapeutic enzymes even when only little labelled training data is available. All four tested enzymes showed high activity, in particular, K3 showed 2-fold better enzymatic activity than the published best KYNase, Pf-K and stronger tumor growth inhibition in animal models. Since the KYNase is originated from bacteria, K3 administration could cause immunogenicity risk *in vivo*, although bacterial enzymes, such as asparaginases, have been FDA-approved and used for certain cancer treatment^17^. PEGylation may help shield KYNase from immune recognition^18^; thus PEG-K3 is open to direct development for anticancer drug.

## Online methods

### Homology search

We searched the kynureninase protein sequence of *Pseudomonas* (WP_017531066.1) in a database of cultivated^8^ and metagenomic protein sequences from BFD^9^ and Metaclust2^10^, a total of 3,144,354,589 protein sequences, using MMseqs2’s (version 3fa46) iterative search (--num-iterations 3 -s 7.5 -c 0.9). The search resulted in 10,195 hits with an E-value of less than 10^-3^ and a bi-directional length overlap of at least 90% similarity. To reduce redundancy, we filtered out sequences with at least 90% sequence identity to another sequence in the set using the filterresult module of MMseqs2, resulting in 5,690 remaining sequences. Since the metagenomic sequences do not contain taxonomic labels, we predicted labels using the taxonomy workflow of MMseqs2^6^ by querying the detected sequences against the UniProtKB/TrEMBL+Swiss-Prot (2020_05) database (As shown in Fig. 1).

### pLLM based regressor to predict *k*_cat_/*K*_M_

To predict the kinetic parameter *k*_cat_/*K*_M_ for enzymes, we followed the variant prediction workflow proposed in the ESM repository (“examples/sup_variant_prediction.ipynb”). Utilizing the ESM-1b model with 670M parameters pretrained on UniRef50, we collected 159 protein sequences containing measurements of *k*_cat_/*K*_M._ Seven originate from the previously published measurements of sequences from *Homo sapiens, Mus musculus, Mucilaginibacter Paludis, Acinetobacter Baumannii, Cyclobacterium marinum, Chlamydophila pecorum* and *Pseudomonas fluorescens*^*5*^. The remaining 152 are modified *H. sapiens* sequences with an average 96.5% sequence identity to the reference, extracted from the patent^11^. For each sequence, we generated ESM-1b sequence embeddings by averaging all residue representations across the whole protein, converting the amino acid sequences into numerical vectors of size 1280.

Subsequently, the dataset was randomly split into training and test sets (80% training, 20% test). We trained three regression models to predict the kinetic parameter, including K-Nearest Neighbors (KNN), Support Vector Regression (SVR), and Random Forest Regression (RFR). Hyperparameter tuning was conducted using GridSearchCV from scikit-learn, optimizing for the best predictive performance based on the R^2^ metric.

Finally, each model’s performance was evaluated on the test dataset by predicting *k*_cat_/*K*_M_. The Spearman rank correlation coefficient was calculated to assess the models’ predictive performance, yielding 0.802, 0.756, and 0.813 for KNN, SVR, and RFR, respectively. Thus, we decided to use the random forest model to rank the proteins detected by the homology search. As a validity check, our RFR model predicted and ranked the enzyme efficiency for all 19 non-wild-type mutations of *H. sapiens* sequence. Notably, 56 of the top 100 mutations were near the enzyme’s active site (position 240–290), confirming the model’s accuracy in identifying key functional regions.

### Webserver as Colab Notebook

We’ve developed a user-friendly Jupyter Notebook, accessible via Google Colaboratory, designed for training customized prediction models using protein sequences and experimental data provided by users in FASTA format. Users input protein sequences in FASTA format with experimental measure. These sequences are transformed into embeddings using the ESM-1b model. For prediction, the tool utilizes KNN, SVM, and RFR regressors. Here, in contrast to the method above, we optimize the embeddings by checking the predictive performance for each transformer block using a five-fold cross-validation. Post-optimization, we use the ColabFold webservice^10^, which is a subset of the 3.2B proteins, to identify homologous proteins encodes them into the optimal layer’s embeddings, and then predicts the defined metric, outputting a ranked list of proteins based on these predictions.

### Plasmid construction

The putative kynureninase and previously reported kynureninase genes from *Pseudomonas fluorescens* were synthesized after codon optimization for heterologous expression in *E. coli* (Gene Universal). Then, the synthesized gene fragments were inserted into the pET-28b(+) vector using NdeI and XhoI as the cut sites to express N-His_6_-tagged kynureninases (Supplementary Table 1).

### Preparation of kynureninases

The custom-ordered plasmids were transformed into *E. coli* BL21(DE3) cells (New England Biolabs). After overnight growth of transformed cells in LB media, the culture (10 mL) was added to TB media (1 L) containing 100 μg/ml of kanamycin in a 4 L-baffled flask. The culture continued at 37 °C in an orbital shaker (170 rpm) until the OD_600_ reached 1.0. Before the addition of isopropyl β–*D*-1-thiogalactopyranoside (IPTG) to the final concentrations of 200 μM, the culture was cooled in an ice bath for 10 min. Then, the culture was shaken at the rate of 170 rpm for 16 h at 15 °C. Cells were harvested by centrifugation (5,000 rpm, 10 min, 4 °C), and the cell pellet was stored at −80 °C until cell lysis. The cell pellet was thawed in Dulbecco’s phosphate buffered saline (DPBS buffer from Alfa Aesar) containing 1 mM pyridoxal phosphate (PLP from Tokyo Chemical Industry, cofactor) at 4 °C, and lysed by microfluidizer (20,000 psi, 2 cycles, LM20 Microfluidics Inc.). After centrifugation (13,000 rpm, 45 min, 4 °C) to remove cell debris, the supernatant was filtered through a 0.45 μm syringe filter and loaded to a pre-equilibrated HisTrap HP column (Cytiva) at 4 °C using ÄKTA Pure FPLC system (Cytiva). After lysate loading, the column was washed with 8 column volume (CV) of DPBS and the protein was eluted by applying the DPBS containing 300 mM imidazole. The fractions colored in yellow were collected (Supplementary Fig. 2) and concentrated using Amicon centrifugal filters (30 kDa cutoff) at 4 °C. After the incubation with PLP (1 mM final concentration) at 25 °C for 1 h, the protein was applied to size exclusion chromatography (HiLoad Superdex 200 pg column, Cytiva) using DPBS at 4 °C. The purity of protein in each fraction was determined using SDS-PAGE (Supplementary Fig. 2). For long-term storage, glycerol-containing buffer (20% v/v) was used and stored at −80 °C until further use.

### Determination of kinetic parameters

The steady-state specific activity assays were carried out, as reported previously^5,12^. In short, we mixed 20 µL of enzyme in DPBS buffer with 80 µL of L-KYN (Sigma-Aldrich) at 37 °C. The consumption rate of substrate was monitored at 365 nm using a microplate reader (Synergy H1, Biotek) at 37 °C. By applying various concentrations of the substrate (0–700 µM), initial rates were measured. Background reaction rates at high concentration of substrates were subtracted from the observed rates. The kinetic parameters, *k*_cat_ and *k*_cat_/*K*_M_, of enzymes were estimated by fitting into the Michaelis-Menten equation using Origin software (Fig. 2b, Supplementary Fig. 3, and Supplementary Table 2).

### System preparation for MD simulations

All calculations were conducted using the X-ray structures of K0 (PDB code 3E9K) and a bacterial KYNase from *Pseudomonas fluorescens* (Pf-K^*^, PDB code 1QZ9), similar to Pf-K. The missing residues in the original PDB files were modeled with the MODELLER program^19^. The PyMOL program was utilized to build the symmetric protein conformation. Molecular docking of L-KYN were conducted using GalaxyDock3^20^. For ligand docking, the substrate binding site was assigned based on the inhibitor site in the X-ray structures. For each enzyme, the initial complex structure for MD simulation had the lowest docking energy among the conformations where the distance between the amine group of substrates and the internal aldimine linkage (C4A) of PLP is less than 3.5 Å. This selection was crucial as the enzyme reaction initiates with the transamination of the C4A atom of PLP and K247 (amine). The 3D-structure of K3 was modelled with AlphaFold2^21^. The energy-minimized structure of Pf-K^*^/L-KYN served as a template for Alphafold2 modeling.

### Molecular Dynamics Simulations

All-atom molecular dynamics (MD) simulations were performed using the *PMEMD*.*cuda* module of AMBER 20 package^13^ with employing the ff14SB force field^22^. Each of three enzyme-substrate combinations, K0/L-KYN, Pf-K*/L-KYN, and K3/L-KYN was solvated in a cubic box using TIP3P water molecules^23^. Counter ions, Na^+^ and Cl^-^, were added to neutralize with a concentration of 150 mM. The system underwent energy minimization, including 500 steps of steepest descent minimization followed by 500 steps of conjugate gradient minimization. This was followed by the second energy minimization, consisting of 1,000 steps of steepest descent minimization followed by 1,500 steps of conjugate gradient minimization. The equilibration process involved three steps in two phases. In the first equilibration step, the system was heated from 0 K to 310 K over 100 ps with weak harmonic restraints of 10.0 kcal/mol/Å^2^ applied to entire simulation system in the NVT ensemble. The second equilibration step applied the same amount of harmonic restraints to the backbone atoms of the enzyme-substrate complexes for 100 ps. In the third equilibration step, 1.0 kcal/mol/Å^2^ of harmonic restraints were applied to Cα atoms of the complex system for 100 ps. The final equilibration phase was performed in the NPT ensemble by Langevin thermostat^24^ and Berendsen barostat^25^ for 4 ns without restraints. Subsequently, a production simulation was conducted for 200 ns, which was repeated 10 times for each system. A cutoff of 10 Å was employed for non-bonded interactions. Long-range electrostatic forces were calculated with the Particle Mesh Ewald method^26^.

### Trajectory Analysis

The MD trajectories were analyzed using *CPPTRAJ*^27^ and in-house Python scripts. An atomic distance fluctuation was calculated using the distances between N of aniline ring of L-KYN and side-chain polar O of binding pocket residues. For hydrophobic residues such as Phe, CB atom was selected. Each snapshot was taken every 1 ns for 200 ns simulation. Cavity volume analysis was performed by CAVER 3.0 built in PyMol^28^.

### pH-dependent activity assay

Specific activities were determined by measuring consumption rates of L-KYN (700 µM) at 25 °C in pH 5.0–8.0 buffers; 50 mM citric acid (pH 5.0–6.0), and potassium phosphate (pH 6.5–8.0). Before measuring the specific activity, enzymes were pre-incubated in each buffer for 10 min at 25 °C.

### Preparation of PEGylated kynureninases

The recombinant proteins were expressed and purified as described above, and phase separation method was applied to remove endotoxin as described previously^29^. In short, Triton™ X-114 (Sigma-Aldrich, 1% v/v) was directly added to the protein solution and mixed by pipetting at 4 °C. After incubation for 10 min, the samples were warmed at 37 °C for 5 min to allow phase separation. Samples were then centrifuged (13,000 rpm, 1 min, 37 °C), and the upper yellow aqueous phase was collected using a micropipette. This procedure was repeated twice to ensure that low endotoxin level. Before PEGylation, the resulting samples were incubated with 1 mM PLP at 25 °C for 1 h. Then, 100-fold molar excess methoxy-PEG-CO(CH_2_)_2_COO-NHS to the protein (5 kDa, NOF America Corporation) was directly added as powder and incubated for 1 h at 25 °C. The unreacted PEGylation reagents were removed by centrifugal filters at 4 °C. The endotoxin level was determined using a chromogenic endotoxin quantification kit (*Limulus* amebocyte lysate from Thermo Fisher Scientific). The purity and the extent of PEGylation were determined using SDS-PAGE.

### Thermal stability of kynureninase activity

Non-PEGylated and PEGylated proteins (10 µM) were incubated at 37 °C using thermal cycler (Bio-Rad Laboratories). After 12 and 24 h, specific activities were measured in the presence of L-KYN (700 µM) in DPBS at 37 °C as described above. Alternatively, we measured the CD spectra of KYNases under various temperatures.

The CD spectra were recorded within the far-UV range (190–250 nm) at a protein concentration of 1 µM in 10 mM sodium phosphate buffer, pH 7.4 at 25 °C, in a 1 mm path length quartz cuvette (Jasco) using a spectropolarimeter (J-815, Jasco). For each sample, ten accumulated scans were averaged and then converted into mean residue ellipticity. The thermal denaturation of non-PEGylated and PEGylated proteins (5 µM) was measured by following the change in ellipticity at 222 nm as a function of temperature increasing from 25 to 95 °C. The melting temperatures were determined by fit the data to the equation of a sigmoidal function.

### Cell lines and mice

The mouse B16-F10 melanoma and CT26 colon cancer cell lines were purchased from the ATCC (American Type Culture Collection, Manassas, Virginia, USA, #CRL-6475). Cell line was maintained in DMEM (Dulbecco’s Modified Eagle’s Medium, Corning, New-York, USA #10-013-CV) with 10% FBS (Fetal Bovine Serum, Corning, New-York, USA #35-015-CV) and 1% penicillin/streptomycin (Thermo, Waltham, MA, USA, #10378016) under the condition of 37 °C in a 5% CO_2_ humified incubator. C57BL/6 inbred mice were purchased from Jackson lab (Bar Harbor, Maine, USA) and BALB/c mice were from Raonbio (Yongin, South Korea). In-vivo studies were approved by the Institutional Animal Care and Use Committee (IACUC, #2022-12-098) of Asan hospital research institute and that of HELIXITH animal research center (VIC-22-06-004) and performed following the approved procedure.

### *In vitro* T cell activation

CD8^+^ T cells were purified from spleen of C57BL/6J mouse using EasySepTM Mouse CD8^+^ T cell isolation kit (STEMCELL, Vancouver, Canada, #19853) by using an EasySep™ magnet (STEMCELL, Vancouver, Canada, #18000) following the protocols provided by the supplier. Purified CD 8^+^ T Cells were plated on a round bottom 96well plate precoated with 5μg/mL of mAnti-CD3 (clone:145-2C11, ebioscienceTM, CA, USA, #16-0031-96) in a T cell activation media consisting of RPMI1640 (#22400-071), 10% FBS (#26140-087), β-mercaptoethanol (55μM, #21985-023), 1X GlutaMax (#35050-061), 100 mM penicillin/streptomycin (#10378-016) and 200 mM sodium pyruvate(#11360-070), that purchased from Gibco (Waltham, Massachusetts, USA), mAnti-CD28 (clone:37.51, ebioscienceTM, CA, USA, #16-0281-86, 2μg/mL) and rhIL-2 (PEPROTECH, NJ, Cranbury, USA, #200-02, 400U/mL), in the presence or absence of L-KYN (Sigma, MA, USA, #K8625-100. 1mM) and of Pf-K-PEG or K3-PEG (0.02uM or 0.05μM). The aggregated T cell group size, indicative of T cell proliferation level, was observed under the microscope (ZEISS primovert, Oberkochen, Germany, #491206-0001-000) on day 5 after activation. Additionally, The cell number was counted with hemocytomter (MARIENFELD, Lauda-Königshofen, Germany, # 0640010) on day 7 after activation.

### Syngeneic implant tumor model

To establish a syngeneic implant tumor model, B16-F10 melanoma and CT26 colon cancer cell lines were implanted subcutaneously into the flank of female C57BL/6 mice (6 wks) and BALB/c mice, respectively. When the tumor volume reaches about 50mm^3^(on 7days post injection), mice were classified equally and randomly into 3 groups (n=13). Pf-K-PEG [dosage:20 mg/kg] and K3-PEG [dosage:20 mg/kg] via peritumoral injections were administered to the mice a total of three times twice a week. The tumor volume was measured twice a week, calculated according to the common formula : V = (Length x Width^2^) x 1/2, T/C ratio(%) was measured as the tumor volume in the control group to the tumor volume in the treated group. TGI(%) was measured as the tumor volume in the control group to the tumor volume in the treated group according to formula : (V_c_ – V_t_)/ V_c_ x 100, Vc = Tumor volume in the control group at the time of tumor extraction / end point. Vt = Tumor volume in the treatment group at the time of tumor extraction / end point.

## Data availably

We provide the data used to perform the analysis at 10.5281/zenodo.10517668. We also build an executable version of the workflow, which is available at seekrank.steineggerlab.com.

## Authors’ Contributions

C.S., M.S.L., W.J.S., and M.S. designed and supervised the research, S.K, M.S., S.P. developed code and performed bioinformatic analyses, H.K. design and conceptualized the figures, Jihyeon.L., Juyong.L. performed MD analysis with the contribution by J.Y., H.E. performed chemical assays and analysis with the contribution by Y-H.H., K.S.C. performed biological assays and analysis, H.E., M.S.L., W.J.S., and M.S. wrote the manuscript.

## Acknowledgment

We thank Milot Mirdita for discussion and helping to revise the manuscript.

## References

1. Dhar, R. et al. Cancer immunotherapy: Recent advances and challenges. J. Cancer Res. Ther. 17, 834–844 (2021).

2. Tang, T. et al. Advantages of targeting the tumor immune microenvironment over blocking immune checkpoint in cancer immunotherapy. Signal Transduct. Target. Ther. 6, 72 (2021).

3. Van den Eynde, B.J., van Baren, N. & Baurain, J.-F. Is there a clinical future for IDO1 inhibitors after the failure of epacadostat in melanoma? Annu. Rev. Cancer Biol. 4, 241–256 (2020).

4. Cheong, J.E. & Sun, L. Targeting the IDO1/TDO2–KYN–AhR pathway for cancer immunotherapy–challenges and opportunities. Trends Pharmacol. Sci. 39, 307–325 (2018).

5. Triplett, T.A. et al. Reversal of indoleamine 2,3-dioxygenase-mediated cancer immune suppression by systemic kynurenine depletion with a therapeutic enzyme. Nat. Biotechnol. 36, 758–764 (2018).

6. Mirdita, M., Steinegger, M., Breitwieser, F., Söding, J. & Levy Karin, E. Fast and sensitive taxonomic assignment to metagenomic contigs. Bioinformatics 37, 3029–3031 (2021).

7. Rives, A. et al. Biological structure and function emerge from scaling unsupervised learning to 250 million protein sequences. Proc. Natl. Acad. Sci. U.S.A. 118, e2016239118 (2021).

8. Consortium, U. UniProt: a worldwide hub of protein knowledge. Nucleic Acids Res. 47, D506–D515 (2019).

9. Steinegger, M., Mirdita, M. & Söding, J. Protein-level assembly increases protein sequence recovery from metagenomic samples manyfold. Nat. Methods 16, 603–606 (2019).

10. Mirdita, M. et al. ColabFold: making protein folding accessible to all. Nat. Methods, 1–4 (2022).

11. Georgiou, G., Stone, E., Blazeck, J., Karamitros, C. Human kynureninase enzymes and uses thereof. US patent 2019/0350975 A1 (2019).

12. Karamitros, C.S. et al. Conformational dynamics contribute to substrate selectivity and catalysis in human kynureninase. ACS Chem. Biol. 15, 3159–3166 (2020).

13. Case, D.A. et al. Amber 2021. (University of California, San Francisco, 2021).

14. Vaupel, P., Kallinowski, F. & Okunieff, P. Blood flow, oxygen and nutrient supply, and metabolic microenvironment of human tumors: a review. Cancer Res. 49, 6449–6465 (1989).

15. Long, G.V. et al. Epacadostat plus pembrolizumab versus placebo plus pembrolizumab in patients with unresectable or metastatic melanoma (ECHO-301/KEYNOTE-252): a phase 3, randomised, double-blind study. Lancet Oncol. 20, 1083–1097 (2019).

16. Hamid, A.B. & Petreaca, R.C. Secondary resistant mutations to small molecule inhibitors in cancer cells. Cancers 12, 927 (2020).

17. Shrivastava, A. et al. Recent developments in L-asparaginase discovery and its potential as anticancer agent. Crit. Rev. Oncol./Hematol. 100, 1–10 (2016).

18. Becicka, W.M. et al. The effect of PEGylation on the efficacy and uptake of an immunostimulatory nanoparticle in the tumor immune microenvironment. Nanoscale Adv. 3, 4961–4972 (2021).

19. Webb, B. & Sali, A. Comparative protein structure modeling using MODELLER. Curr. Protoc. Bioinformatics 54, 5.6.1-5.6. 37 (2016).

20. Yang, J., Baek, M. & Seok, C. GalaxyDock3: Protein–ligand docking that considers the full ligand conformational flexibility. J. Comput. Chem. 40, 2739–2748 (2019).

21. Jumper, J. et al. Highly accurate protein structure prediction with AlphaFold. Nature 596, 583–589 (2021).

22. Maier, J.A. et al. ff14SB: improving the accuracy of protein side chain and backbone parameters from ff99SB. J. Chem. Theory Comput. 11, 3696–3713 (2015).

23. Jorgensen, W.L., Chandrasekhar, J., Madura, J.D., Impey, R.W. & Klein, M.L. Comparison of simple potential functions for simulating liquid water. J. Chem. Phys. 79, 926–935 (1983).

24. Pastor, R.W., Brooks, B.R. & Szabo, A. An analysis of the accuracy of Langevin and molecular dynamics algorithms. Mol. Phys. 65, 1409–1419 (1988).

25. Berendsen, H.J., Postma, J.v., Van Gunsteren, W.F., DiNola, A. & Haak, J.R. Molecular dynamics with coupling to an external bath. J. Chem. Phys. 81, 3684–3690 (1984).

26. Darden, T., York, D. & Pedersen, L. Particle mesh Ewald: An N· log (N) method for Ewald sums in large systems. J. Chem. Phys. 98, 10089–10092 (1993).

27. Spranger, S. et al. Up-regulation of PD-L1, IDO, and Tregs in the melanoma tumor microenvironment is driven by CD8+ T cells. Sci. Transl. Med. 5, 200ra116–200ra116 (2013).

28. Pavelka, A. et al. CAVER: algorithms for analyzing dynamics of tunnels in macromolecules. IEEE/ACM transactions on computational biology and bioinformatics 13, 505–517 (2015).

29. Yari, M., Ghoshoon, M.B., Vakili, B. & Ghasemi, Y. Therapeutic enzymes: applications and approaches to pharmacological improvement. Curr. Pharm. Biotechnol. 18, 531–540 (2017).

