## Supplementary Information for "Discovery of Highly Active Kynureninases for Cancer Immunotherapy through Protein Language Model"

**Table of contents**

**Supplementary Figures**

Supplementary Figure 1. Taxonomic distribution of hits detected by search through UniProtKB and metagenomic databases.

Supplementary Figure 2. Purification of KYNases.

Supplementary Figure 3. Measured steady-state kinetic parameters.

Supplementary Figure 4. Time-dependent consumption of L-KYN by Pf-K and K3.

Supplementary Figure 5. The time-series of inter-atomic distances between the amine nitrogen of aniline ring of L-KYN and three key residues forming the active site in Pf-K and K3.

Supplementary Figure 6. The time-series of inter-atomic distances between the amine nitrogen of aniline ring of L-KYN and three residues from the active site of each monomer of K0.

Supplementary Figure 7. pH-dependent activity of Pf-K and K3.

Supplementary Figure 8. CD spectra of non-PEGylated and PEGylated Pf-K and K3.

Supplementary Figure 9. Reversal of L-KYN-induced inhibition of T cell activation/proliferation *In vitro*.

Supplementary Figure 10. Antitumor effects of Pf-K-PEG and K3-PEG *in vivo*.

Supplementary Figure 11. The change of body weight during the PEG-KYNase treatment.

Supplementary Figure 12**.** Antitumor effect of PEG-KYNase in colon cancer mouse model (CT26).

**Supplementary Tables**

Supplementary Table 1. The amino acid sequences of kynureninases used in this study.

Supplementary Table 2. Steady-state kinetic parameters of kynureninases used in this study.

Supplementary Table 3. Cavity volume in the active site of Pf-K and K3.

**Supplementary Figure 1.** Taxonomic distribution of hits detected by search through UniProtKB and metagenomic databases. Taxon were predicted by MMseqs2 taxonomy. Sankey plot was produced by Pavian^1^.

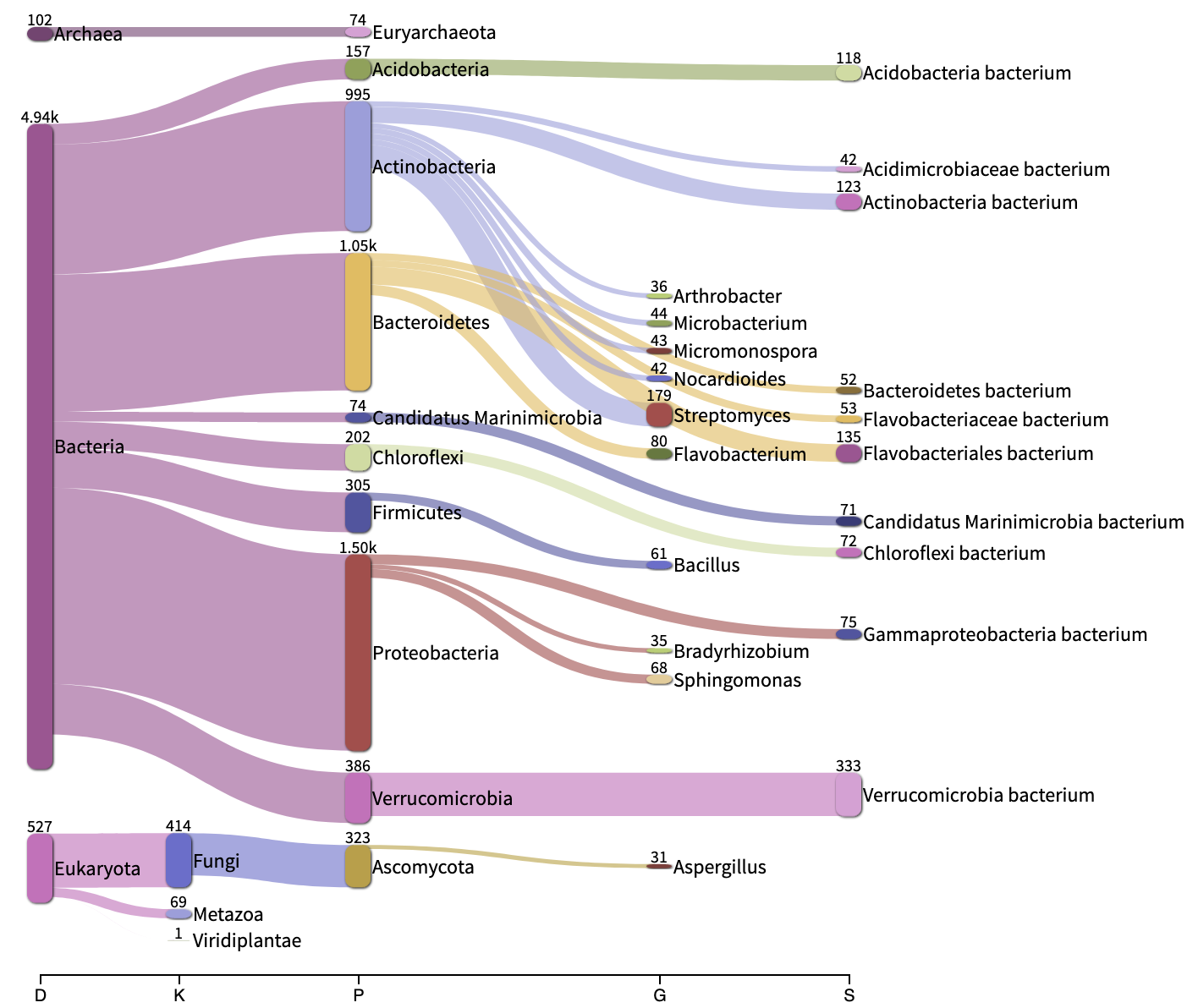

**Supplementary Figure 2.** Purification of KYNases. An elution profile of (**a**) HisTrap affinity chromatography and (**b**) size-exclusion chromatography. The elution of kynureninase is marked with an asterisk. (**c**) SDS-PAGE of K1-K31. (**d**) Representative UV-Vis absorption spectrum of Pf-K. The absorption at 430 nm region indicates the presence of PLP-cofactor.

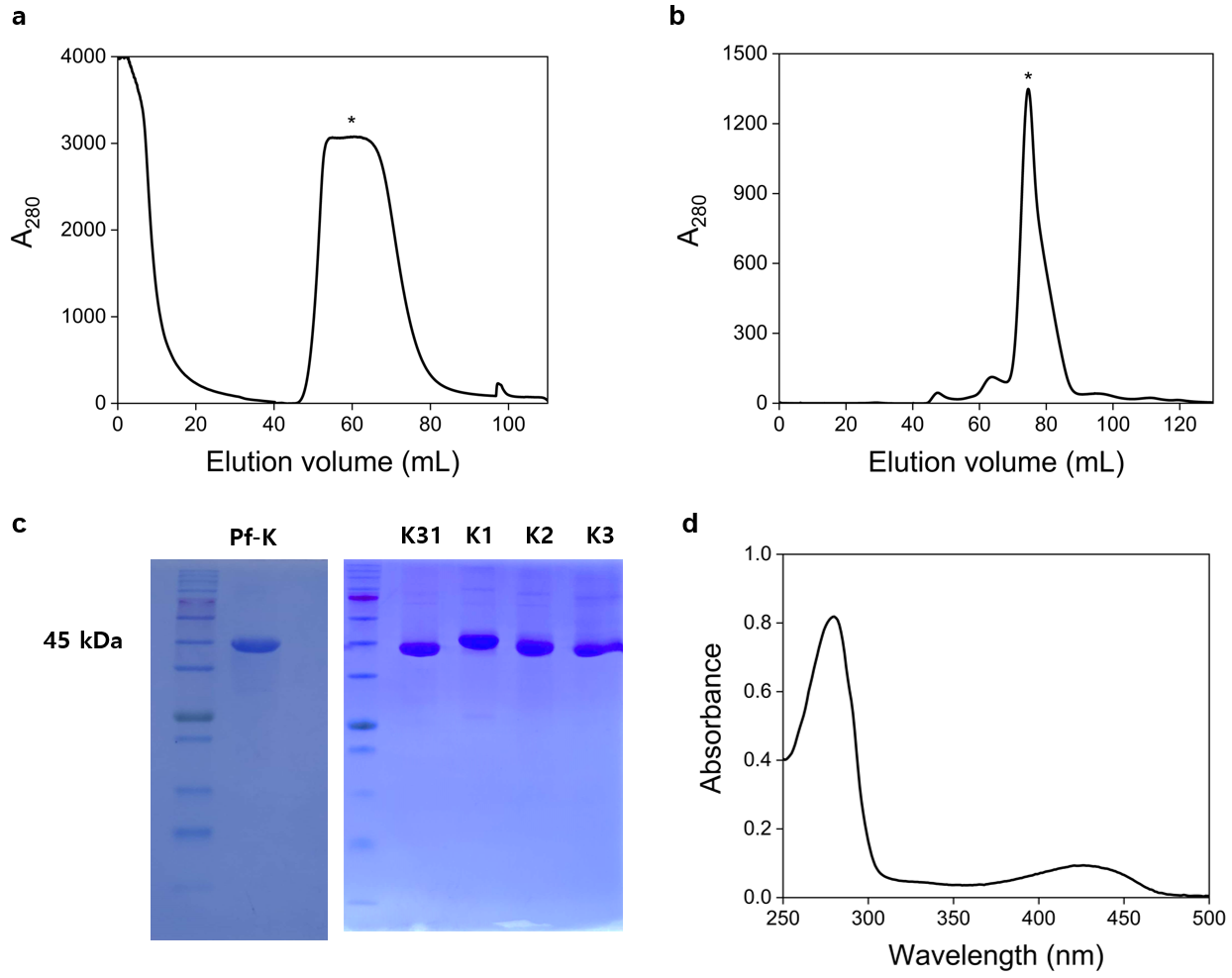

**Supplementary Figure 3.** Measured steady-state kinetic parameters. (**a**) *k*_cat_ values and (**b**) *k*_cat_/*K*_M_ values for KYNases. Error bars indicate the standard error and *p* values from two-sample *t* tests (** *p* ≤ 0.01, ns; not significantly different at the *p* < 0.05 level).

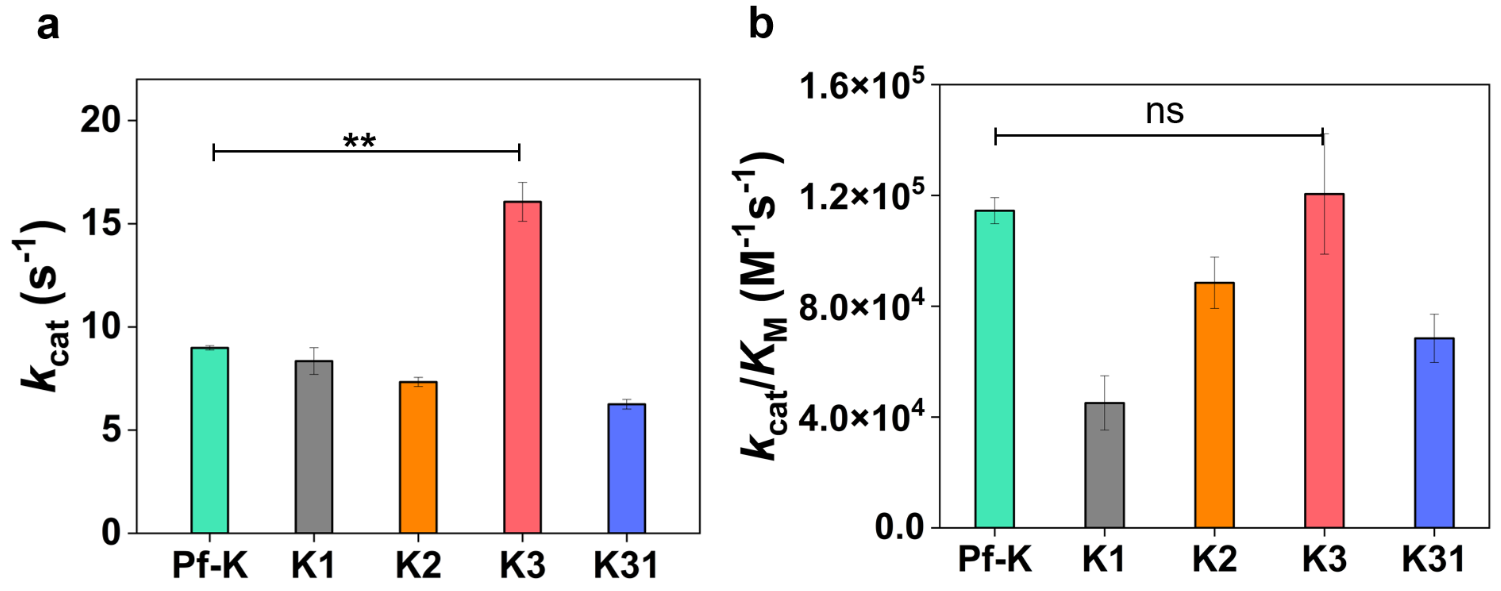

**Supplementary Figure 4.** Time-dependent consumption of L-KYN by Pf-K and K3. Time-dependent consumption of substrate (700 µM) was monitored by measuring absorbance (365 nm) at 37 °C.

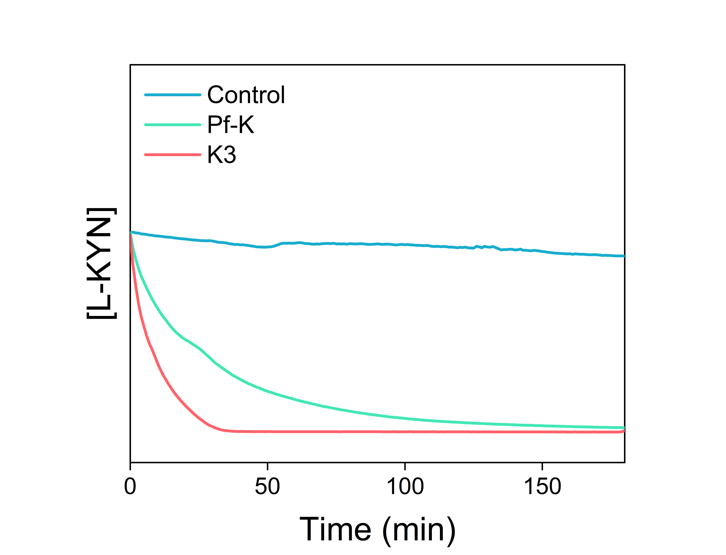

**Supplementary Figure 5.** The time-series of inter-atomic distances between the amine nitrogen of aniline ring of L-KYN and three key residues forming the active site in (**a**) Pf-K and (**b**) K3. Residues marked with asterisks (*) are from the other symmetric monomer.

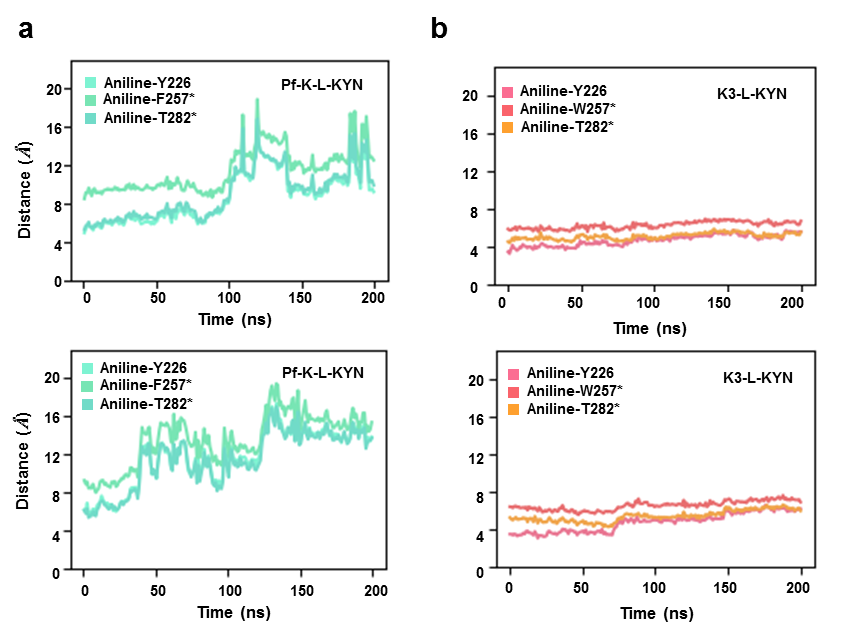

**Supplementary Figure 6. (a)** The time-series of inter-atomic distances between the amine nitrogen of aniline ring of L-KYN and three residues from the active site of each monomer of K0. The asterisk marks (*) denote residues from the other symmetric monomer. (**b**) The dotted yellow line emphasizes L-KYN out of binding site due to high fluctuation.

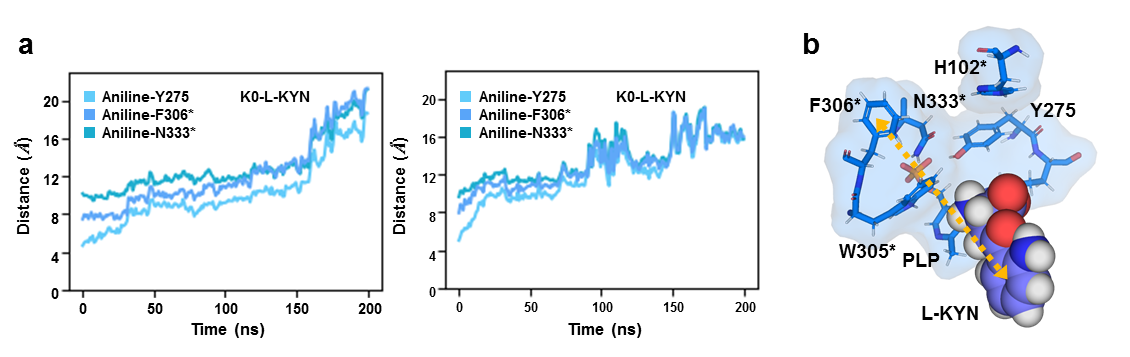

**Supplementary Figure 7**. pH-dependent activity of Pf-K and K3. Specific activities were determined by measuring consumption rates of *L*-kynurenine (700 µM) at 25 °C in pH 5.0–8.0 buffers, 50 mM citric acid (pH 5.0–6.0), and potassium phosphate (pH 6.5–8.0). Error bars indicate the standard error and *p* values from two-sample *t* tests (* *p* ≤ 0.05, ** *p* ≤ 0.01, *** *p* ≤ 0.001).

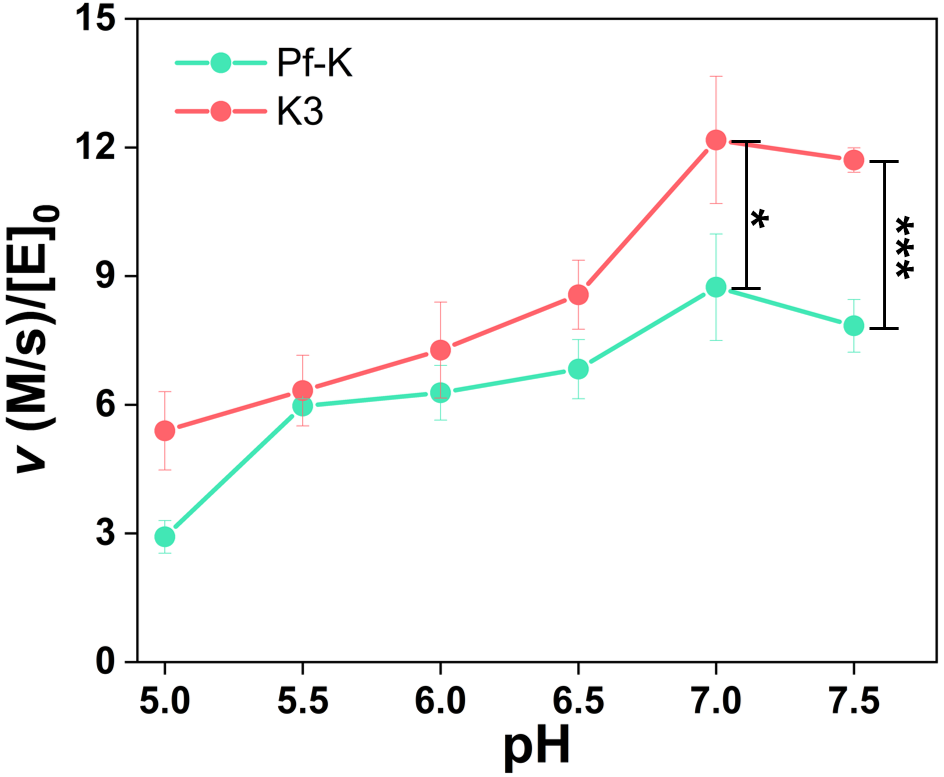

**Supplementary Figure 8.** CD spectra of non-PEGylated and PEGylated Pf-K and K3. (**a**) As isolated forms, the CD spectra were recorded within the far-UV range (190–250 nm) at a protein concentration of 1 µM in 10 mM sodium phosphate buffer, pH 7.4 at 25 °C. Ten accumulated scans were averaged for each sample and then converted into mean residue ellipticity. (**b**) Temperature-dependent CD spectral changes. The thermal denaturation of non-PEGylated and PEGylated proteins (5 µM) was measured by following the change in ellipticity at 222 nm as a function of temperature increasing from 25 to 95 °C.

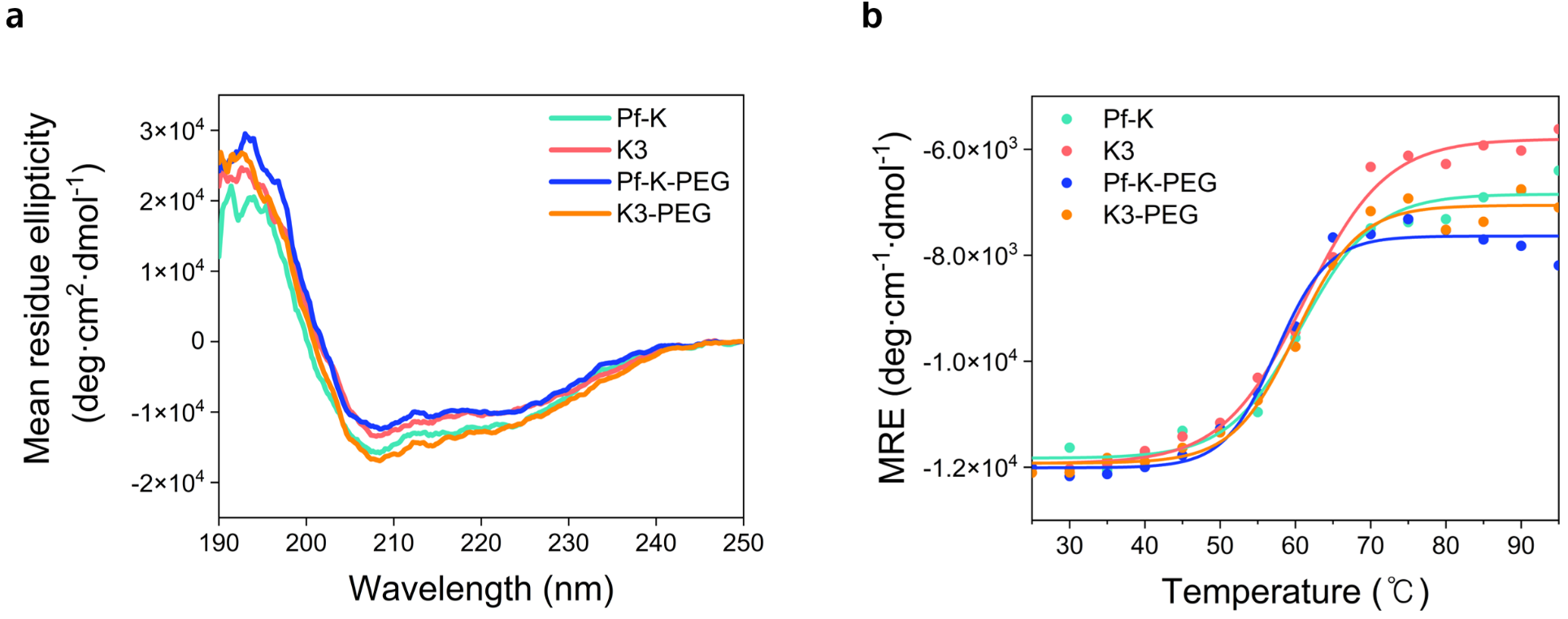

**Supplementary Figure 9.** Reversal of L-KYN-induced inhibition of T cell activation/proliferation *In vitro*. Representative CD8^+^ T cell group images after TCR stimulation for 5 days in the presence of Pf-K-PEG or K3-PEG (0.02 μM or 0.05 μM) and *L*-Kyn (1 mM).

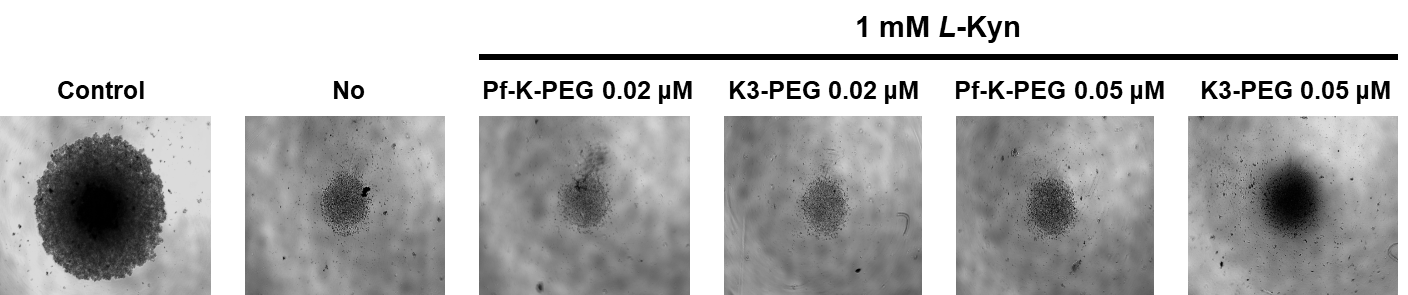

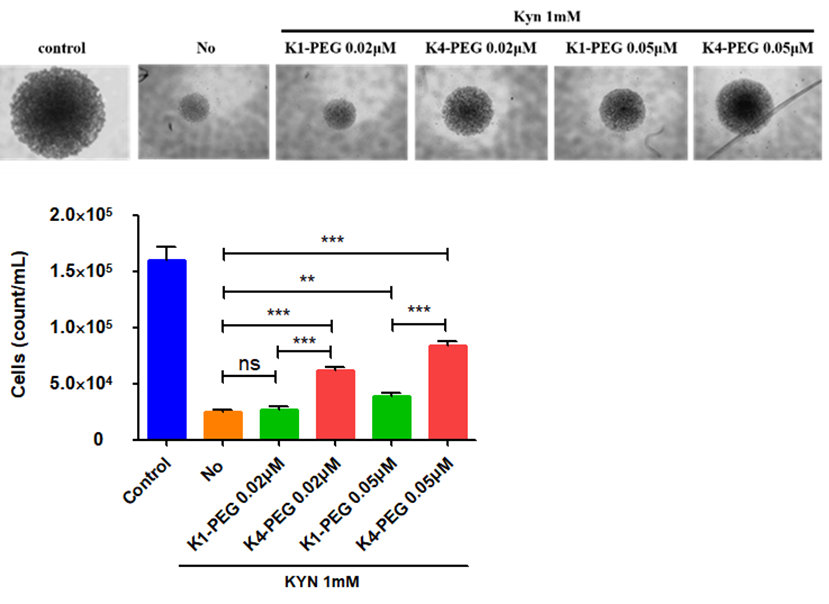

**Supplementary Figure 10.** Antitumor effects of Pf-K-PEG and K3-PEG *in vivo*. Tumor growth curve of B16-F10 bearing mice (C57BL/6) that were treated with peritumoral injection of control (PBS, n=13), Pf-K-PEG (20 mg/kg, n=13) and K3-PEG (20 mg/kg, n=13). Arrows indicate treatment point.

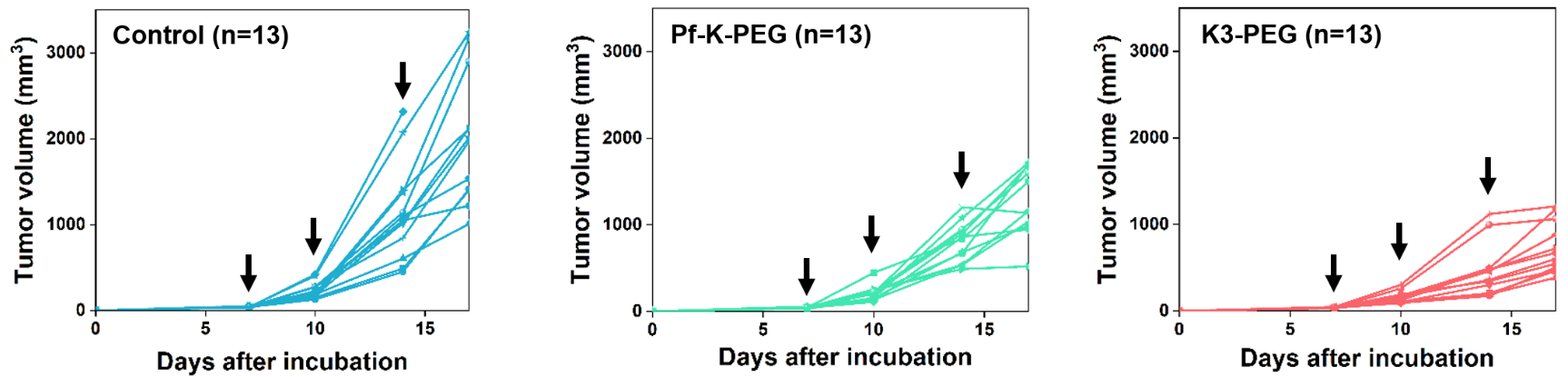

**Supplementary Figure 11.** The change of body weight during the PEG-KYNase treatment. B16-F10 bearing mice (C57BL/6) were treated with a peritumoral injection of control (PBS, n=13), Pf-K-PEG (20mg/kg, n=13) or K3-PEG (20mg/kg, n=13) twice a week (total 3 times) and the body weight was measured twice a week after treatment until the end of the experiment. There is no statistically significant difference between treatment groups.

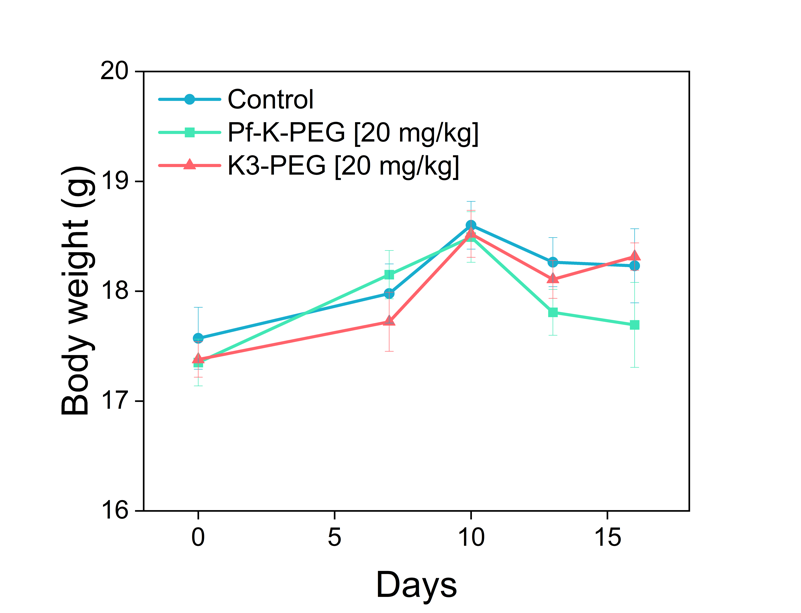

**Supplementary Figure 12.** Antitumor effect of PEG-KYNase in colon cancer mouse model (CT26). Tumor growth curve of CT26 bearing mice (BALB/c) that were treated with a peritumoral administration of 20 mg/kg (vehicle, Pf-K-PEG or K3-PEG). Mean and individual tumor growth curves over time. Statistical significance was verified using a two-tailed t-test (* *p* ≤ 0.05, ** *p* ≤ 0.01, *** *p* ≤ 0.001). Error bars represent the mean± SEM.

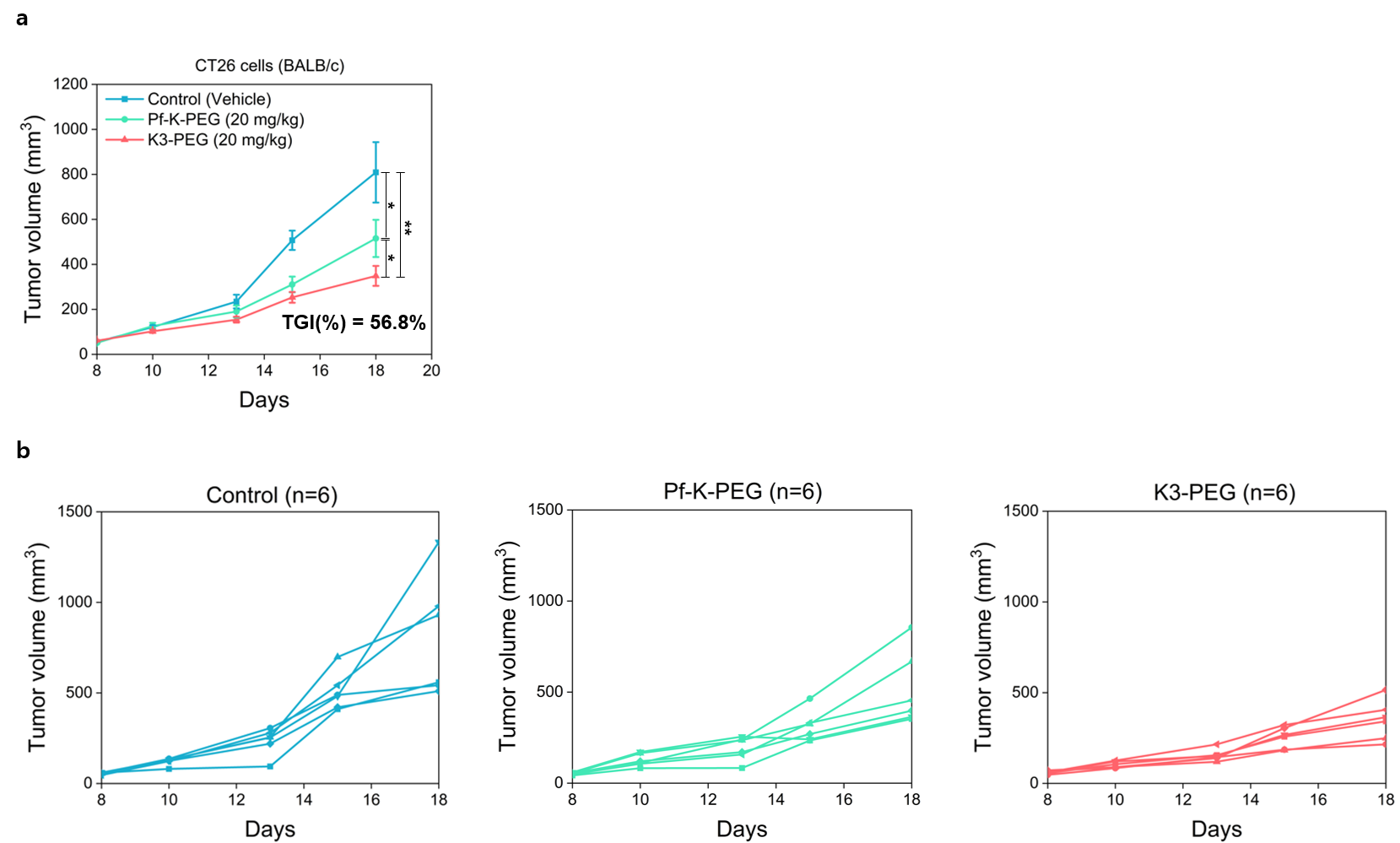

**Supplementary Table 1.** The amino acid sequences of kynureninases used in this study. Amino acid sequences without tags are shown in bold.

| Protein | Amino acid sequence |
| --- | --- |
| Pf-K from *Pseudomonas fluorescence* | MGSSHHHHHHSSGLVPRGSH**MTSRSHCQTLDAQDPLAPLRDQFALPAGVIYLDGNSLGARPVASLARAQQVIAEEWGNGLIRSWNSAGWADLSLRLGNRLAPLIGAGAGEVAITDTTSINLFKVLSAALTVQRQREPARKVIVSEASNFPTDLYIAEGLAELLQQGYCLRLVNSPDELPQAIDADVAVVMLTHVNYKTGYMYDMQALTALSHECGALSIWDLAHSAGAVPIDLRAAGADYAIGCTYKYLNGGPGSQAFVWVNPALVDQVRQPLSGWFGHTRQFAMESNYAPSAGIARYLCGTQPITSLAMVECGLQIFEQTDMACLRRKSLALTDLFIALVEARCAAHGLVLITPREHARRGSHVSFEHPEGYAVIQALIARGVIGDYREPRIMRFGFTPLYTRFSEVWDAVEILGEILDESTWDQPQFKVRHSVT** |
| K1 from *Pseudomonas sp.* | MGSSHHHHHHSSGLVPRGSH**MATQQHLRELDLADPLAALRDEFALPEGVIYLDGNSLGARPKAALARATQMIEQEWGEGLIRSWNTAQWSTLSARLGDKLAPLIGADSGEVVITDTTTVNLFKILAAGLRIQAERAPQRKIILSELHNFPADLYVIEGLADLLQQGYELRLIERPQDLPGLLDDQVALVLLTHVNYKSGHMYDMTATTGLIHQHGALALWDLAHSAGAVPVQLKAANADYAIGCTYKYLNGGPGSPAFVWVSPQLCDQVWQPLSGWWGHARQFAMEPHYAPASGITRYLCGTQPLVSLGMVESGLDIFAKTSMQALRNKSLALTDLFIELVEARCKDHPLTLITPRDHAQRGSHVSFEHPEGYAVVQALIARGVIGDYREPRIMRFGFTPLYTSFQDVGAAVQALVEVLDSQQWREPQFQTRHSVT** |
| K2 from *Pseudomonas sp.* | MGSSHHHHHHSSGLVPRGSH**MITRNDCLALDAQDPLAHLRHQFALPEGVIYLDGNSLGARPIAALERAQAVIAEEWGNGLIRSWNSAGWLDLPERLGNRLAGLIGAGEGEVVVTDTTSINLFKVLGAALRVQAMRAPTRRVIVSESSNFPTDLYIAEGLMDLLQQGYSLRLVDSPEELAQAIDQDTAVVMLTHVNYKTGYMHDMQAVTVLIHECGALAIWDLAHSAGAVPVDLRQAGADYAIGCTYKYLNGGPGSQAFVWVAPQLCDLVTQPLSGWFGHSRQFDMASGYEPSSGIARYLCGTQPITSLAMVECGLEIFAQTDMPSLRRKSLALTDLFIQMVEQRCAAHDLKLITPREHARRGSHVSFEHPQGYAVIQALIARGVIGDYREPRIMRFGFTPLYTSFTEVFDAVQILGEILDQQTWSQAQFQVRHSVT** |
| K3 from *Bordetella sp.* | MGSSHHHHHHSSGLVPRGSH**MHTREACLQADQQDPLAPLKAQFDIPAGVLYMDGNSLGVLPKAAVARSAQVIQQEWGQGLIRSWNDASWFELPSRLGDKLGRLIGAGTGQVVVTDTTSLNLFKSLAAAIRIQQQAAPQRKIIVSERDNFPTDLYMIQGMIDLLQQGYEMRLVDEDLSLEQALDDSVAVLLLSHVNYRTGHMYDMADVTAQAHARGALTIWDLAHAAGAVPVDLTGANADFAVGCTYKYLNGGPGAPAFIWVAPRHTDHFWQPLSGWWGHQRPFDMAVNYEPAGGIRRYLCGTQPIVSLSLVECGLDISLQADMNEVRRKSLALTDLFIALVESRCARHPLTLVTPREHAHRGSHVSLRHPHGYAVMQALIARGVIGDYREPEVLRFGFTPLYFGYTDVWDAVEILTDVLDSEIWKQPEFSRRGAVT** |
| K31 from *Rheinheimera sp.* | MGSSHHHHHHSSGLVPRGSH**MTCAALQQRDIDDPLSGKRAAFYLPDNTLYLDGNSLGAMPKIAAERAAEVVSQQWGEGLITSWNRHHWIDLPFSVGDKIGHLIGAAPGQVICCDSTSVNLFKVLCAALSLQPARSKVLSVSGNFPTDLYMVEGLSALTGNNHYQLQLVDESELEQAITGQVAVLLLTHVDFRSGRLFDMAKLTRLAQDKGALVIWDLAHSAGALPLALDQCHVDFAVGCGYKYLNGGPGAPAFLYAAKRHHAMLQQPLTGWMGHKTPFSFSTQYEKASGIAQFLTGTPPVISMSVLDAALDVFADVDIAQLRQKSLALSDCFHQLVSQNDCLNELERITPYAAAERGSQLAYRHPQAYALCQALIKQGVIADFRAPDILRLGFTPLYLRYIDVWTAVEILADVMRSSEYLKAEYQIKQKVT** |

**Supplementary Table 2.** Steady-state kinetic parameters of kynureninases used in this study. The activities were measured in triplicate.

|  | *Identifier* | *k*_cat_ (s^-1^) | *K*_M_ (μM) | *k*_cat_/*K*_M_ (M^-1^ s^-1^) |
| --- | --- | --- | --- | --- |
| Pf-K | WP_017531066.1 (GenBank) | 9.0(1) | 78(3) | 11(0) $\times$ 10^4^ |
| K1 | A0A0Q5EY38 (Uniprot) | 8.3(6) | 185(38) | 4.5(10) $\times$ 10^4^ |
| K2 | Marine metagenome | 7.3(2) | 83(8) | 8.8(9) $\times$ 10^4^ |
| K3 | A0A261TMR2 (Uniprot) | 16(1) | 133(23) | 12(2) $\times$ 10^4^ |
| K31 | A0A117NJK5 (Uniprot) | 6.2(2) | 91(11) | 6.8(9) $\times$ 10^4^ |

**Supplementary Table 3.** Cavity volume in the active site of Pf-K and K3. KYNases showing the van der Waals interactions with 3-OH-KYN versus L-KYN. The time-averaged values and standard errors for 1,000 snapshots with separate values provided for each symmetry-related monomer 1 and 2.

|  | Monomer 1 [Å^3^] | Monomer 2 [Å^3^] |
| --- | --- | --- |
| Pf-K/3-OH-KYN | 2.4 *±* 0.2 | 2.6 *±* 0.2 |
| Pf-K/L-KYN | 2.4 *±* 0.1 | 2.3 *±* 0.1 |
|  | Monomer 1 [Å^3^] | Monomer 2 [Å^3^] |
| K3/3-OH-KYN | 1.8 *±* 0.1 | 1.8 *±* 0.2 |
| K3/L-KYN | 1.8 *±* 0.1 | 2.0 *±* 0.1 |

**Reference**

1.	Breitwieser, F.P. & Salzberg, S.L. Pavian: interactive analysis of metagenomics data for microbiome studies and pathogen identification. *Bioinformatics* **36**, 1303-1304 (2020).
